# Therapeutic gene editing in hematopoietic progenitor cells from a mouse model of Fanconi anemia

**DOI:** 10.1101/362251

**Authors:** MJ Pino-Barrio, Y Gimenez, M Villanueva, M Hildenbeutel, R Sánchez-Dominguez, S Rodriguez-Perales, R Pujol, J Surrallés, P Rio, T Cathomen, C Mussolino, JA Bueren, S Navarro

**Author notes:** Corresponding authors: Susana Navarro.; Juan A. Bueren.

## Abstract

The promising ability to genetically modify hematopoietic stem and progenitor cells (HSPCs) by precise gene editing remains challenging due to their sensitivity and poor permissiveness. This represents the first evidence of implementing a gene editing strategy in a murine *safe harbor* locus that phenotypically corrects primary cells derived from a mouse model of Fanconi anemia (FA).

By co-delivering TALENs and a donor therapeutic *FANCA* cassette template to the *Mbs85* locus (ortholog of the h*AAVS1 safe harbor* locus), we achieved efficient gene targeting (23%) in FA mouse embryonic fibroblasts (MEFs). This resulted in the phenotypic correction of these cells, as revealed by the improvement of their hypersensitivity to mitomycinC. Moreover, robust evidence of targeted integration was observed in murine WT and FA-A hematopoietic progenitor cells (HPC) reaching mean targeted integration values of 20.98% and 16.33% respectively, with phenotypic correction of FA HPCs. Overall, our results demonstrate the feasibility of implementing a therapeutic targeted integration strategy in a murine *safe harbor* locus, such as the *Mbs85* gene, of MEFs and murine HPC from a FA mouse model.

## INTRODUCTION

Fanconi anemia (FA) is a rare genetic disorder associated with mutations in any of the twenty-two FA genes, known as FANC genes (Bagby, 2018; Knies et al, 2017). The genetic products of these genes belong to a DNA repair pathway known as the FA/BRCA pathway, which is involved in the repair of interstrand cross-link (ICL) lesions during DNA replication. FA patient cells are characterized by the accumulation of DNA damage at an increased rate as compared to healthy cells due to an ineffective FA/BRCA DNA repair pathway. Furthermore, most patients show congenital abnormalities at birth, cancer predisposition (Auerbach, 2009; Schneider et al, 2015; Tischkowitz & Hodgson, 2003), and bone marrow failure (Ceccaldi et al, 2012; Kelly et al, 2007). Due to the risks of allogeneic hematopoietic stem cell transplantation, alternative curative treatments have been proposed. This is the case of gene therapy approaches which aim at the correction of autologous HSPC with therapeutic lentiviral vectors. The efficiency and safety of these strategies in preclinical stages have previously been demonstrated (Becker et al, 2010; Galimi et al, 2002; Gonzalez-Murillo et al, 2010; Jacome et al, 2009; Molina-Estevez et al, 2015; Muller et al, 2008) and are nowadays tested in clinical trials (Adair et al, 2017; Navarro et al, 2015; Tolar et al, 2011; Tolar et al, 2012). However, targeted gene therapy approaches are evolving as a promising alternative to avoid random integration issues arising from the use of retroviral vectors.

Due to the fact that the most frequent complementation group of FA patients is FA-A (60-70%), which is characterized by highly heterogeneous mutations (Casado et al, 2007; Castella et al, 2011) in the *FANCA* gene (Mehta & Tolar, 1993), a therapeutic strategy to precisely integrate a therapeutic *FANCA* expression cassette into a *safe harbor* locus (Papapetrou & Schambach, 2016; Sadelain et al, 2011) would be applicable to all *FANCA* mutations (Diez et al, 2017; Rio et al, 2014) and could be in principle extended to other FA subtypes.

The human *AAVS1* locus located on the first intron of the *MBS85* (*PPP1R12C*) gene on chromosome 19 (Kotin et al, 1992) meets the requirements of a *safe harbor* locus in a wide range of cell types (DeKelver et al, 2010; Hockemeyer et al, 2009; Lombardo et al, 2011; Oceguera-Yanez et al; Ramachandra et al, 2011; Smith et al, 2008; Zou et al, 2011). It has an open chromatin status and also contains a putative insulator element (van Rensburg et al, 2012). Thus, based on the favorable results observed in human cells, we propose to explore a similar strategy in the context of a mouse model of FA-A by integrating a therapeutic *FANCA* expression cassette in the murine *Mbs85 gene*, the ortholog of the human *AAVS1* (Dutheil et al, 2004; Henckaerts & Linden, 2010; Linden et al, 1996a; Linden et al, 1996b). This gene spans 20 kilobases (kb), and the resulting 3.1 kb mouse cDNA and protein are 77% and 86% identical to their human counterparts, respectively (Tan et al, 2001).

Our study demonstrates the feasibility of conducting a targeted gene therapy approach in embryonic fibroblasts and hematopoietic progenitors from a mouse model of FA-A and highlights the potential of using for the first time the *Mbs85* locus as a murine *safe harbor* for targeted integration, what opens a new platform that allows the study of the real implication of what means a safe harbor locus in an *in vivo* model prior to the clinic.

## RESULTS

### Targeted genome integration in FA-A mouse embryonic fibroblasts

To establish efficient genome editing at the murine ortholog of the human *AAVS1* gene, we generated a pair of transcription activator-like effector nucleases (TALEN) (Mussolino et al, 2011) targeting the first intron of the murine *Mbs85* gene (**EV1A**). The expression of each TALEN monomer was then evaluated by western blot analyses in HEK-293T cells upon lipofection of the corresponding plasmids using an antibody recognizing the HA-tag (**EV1B**). Mouse embryonic fibroblasts (MEFs) from *Fanca*^-/-^ mice were used to define the optimal conditions for achieving targeted integration at the murine *Mbs85* locus. *Mbs85*-specific TALENs and a donor construct containing a therapeutic human *FANCA* cassette flanked by homologous sequences to the murine *Mbs85* gene (**Fig 1A and B**) were delivered into the FA-A MEFs via nucleofection using different amounts of the *FANCA* therapeutic donor and a fixed dose of TALEN monomers expression plasmids (**Fig 1C)**.

**Fig 1.**
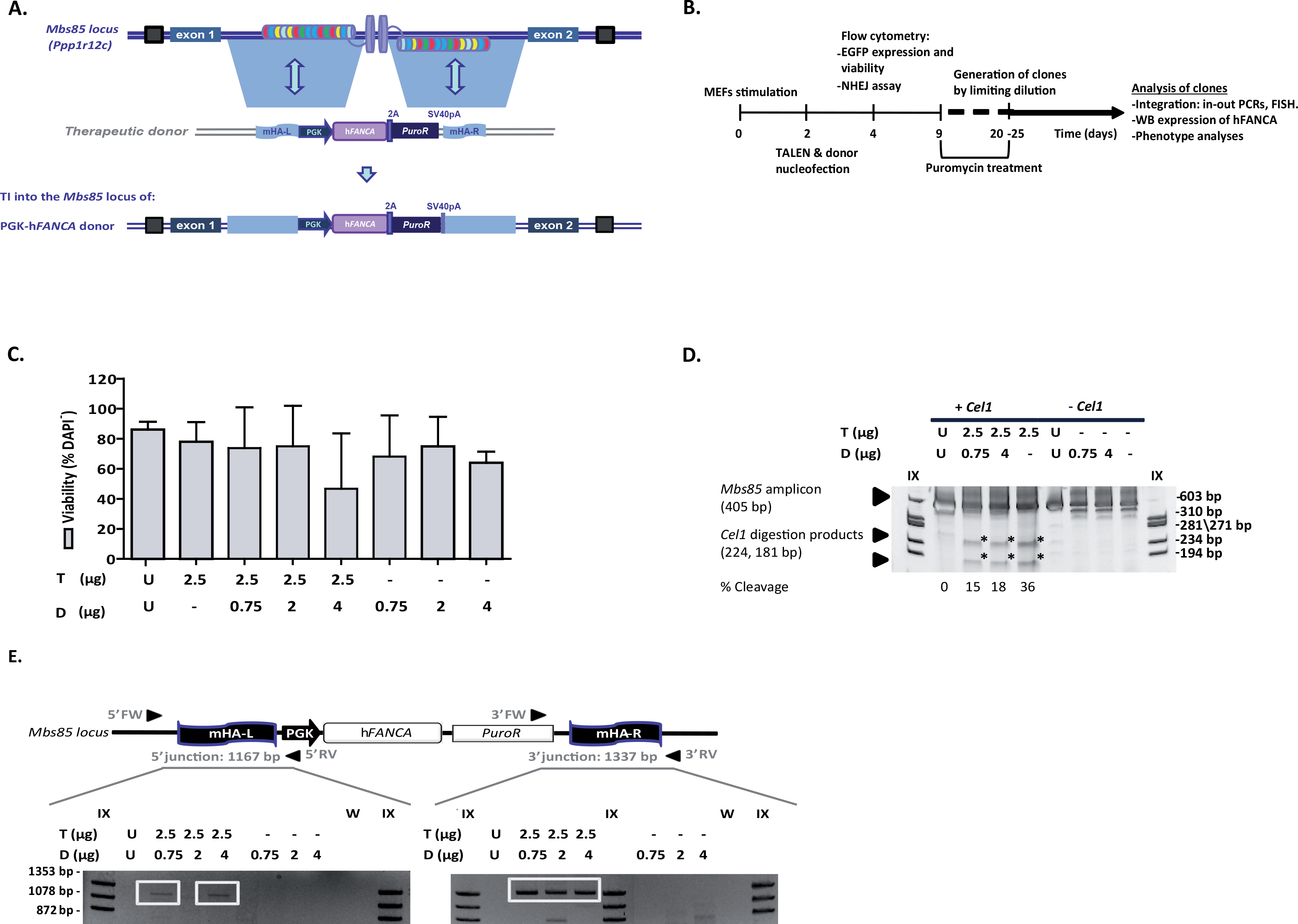
*Mbs85*-specific TALEN and donor-mediated targeted insertion in FA MEFs. **A)** Schematic showing the architecture of the murine *Mbs85* locus, with the target sites of the TALENs highlighted, and the structure of the donor used with the therapeutic h*FANCA* cassette driven by the phosphoglycerate kinase promoter (PGK) flanked by sequences homologous to the genomic target locus. The resulting locus upon targeted integration (TI) of the donor is indicated in the lowest part of the panel. mHA-L and mHA-R: homology arms to the murine *Mbs85* locus; 2A: 2A self-cleaving peptide sequence; SV40pA: simian virus 40 polyA sequence; *PuroR*: Puromycin resistance gene. **B)** Flow chart indicating the study design and the different analyses performed in gene-edited cells of FA-A MEFs. **C)** Analysis of viability (percentage of DAPI^-^ cells) in the different conditions. U: untransfected cells; T: 2.5 µg of each TALEN monomer; D: donor doses (0.75, 2 or 4 µg). Bars indicate the mean ± S.D. (n=2 experiments). **D)** Cleavage efficacy of the *Mbs85*-specific TALENs analysed by Surveyor assay. Representative electrophoresis gel showing the disruption of the target locus in FA-A MEFs nucleofected with only the TALENs (T, 2.5 µg of each TALEN monomer) or together with different donor doses (0.75 µg and 4 µg). U: untransfected cells. Samples not digested with the *Cel1* endonuclease were used as controls. The extent of TALEN cleavage, measured as the mean percentage of modified alleles, is indicated below. Arrows indicate the size of the parental band (405 bp) and the expected positions of the digestion products (224 bp and 181 bp), that are also indicated with asterisks. IX: DNA molecular weight marker. **E)** Schematic representation of the targeted integration of the therapeutic PGK-*hFANCA* donor into the *Mbs85* locus of FA-A MEFs. Arrows represent the primers, forward (Fw) and reverse (Rv) used to evaluate the site-specific integration and the size of the PCR amplicon is indicated for each integration junctions. The electrophoresis gel below is a representative image of the integration analysis performed on the same samples as indicated in C). W: water control; IX: DNA molecular weight marker.

The viability of nucleofected cells was on average around 69% in comparison with 86% of untransfected cells at 48 hours post nucleofection (**Fig 1C**). An *EGFP* control plasmid was nucleofected in these cells as a control to evaluate the transfection efficiency in MEFs, that resulted in 30.17 ± 6.15%. Then, we analyzed the ability of the designed TALEN to disrupt the mouse *Mbs85* locus using Surveyor assay. The frequency of indel mutations ranged from 15% to 36% after nucleofection with these TALENs (**Fig 1D**). The simultaneous delivery of TALENs and donor DNA reduced the frequency of indel mutations by 50%, as compared to delivering the nucleases alone (**Fig 1D**, conditions T+D), suggesting that a number of DSBs were repaired by HR instead of NHEJ.

Once the ability of designed TALENs to cleave the *Mbs85* locus had been corroborated, we evaluated the efficacy of the gene targeting strategy. The presence of a puromycin selection marker in the therapeutic donor allowed us to enrich the population of FA-A MEFs that underwent correct gene targeting. Nucleofected cells were maintained in culture for 5 passages in the presence of puromycin (1-1.25 µg/ml). Diagnostic PCRs were then performed in the bulk population to demonstrate the integration of the therapeutic PGK-h*FANCA* donor into the *Mbs85* locus using the primers indicated to amplify the 5’ and the 3’ integration junctions (**Fig 1E and Supplementary Table S3**). Samples nucleofected with the TALENs and donor revealed successful gene targeting, with amplification of the specific bands corresponding to the insertion of the donor into the *Mbs85* locus (as shown in **Fig 1E**).

A total of 174 clones were generated by limiting dilution with the aim of determining the efficiency of targeted integration in FA-A MEFs. In two independent experiments, a mean of 23.1% of the clones showed site-specific gene targeting in the *Mbs85* locus, reaching a maximum of 7.4% efficiency at the 5’ integration junction. Overall, five clones (nucleofected with 2.5 µg of each TALEN monomer and 0.75 µg of the therapeutic donor) showed the expected 5’ and 3´ integration junctions.

Subsequently, we proceeded to determine the frequencies of insertion/deletion (indel) mutations in the FA-MEFs pool by deep sequencing at the on-target site and the 23 putative off-target sites predicted by PROGNOS (Fine et al, 2013) (**Table1 and Supplementary Table1**). The frequency of TALEN-induced indels at the on-target locus of WT and FA-A MEFs was 37.1%, and 30.1%, respectively. These frequencies were also confirmed by Surveyor assay (**EV1C**). Importantly, the frequencies of indels at three identified off-target sites ranged from 0.06% to 0.36%, showing that the used TALENs were highly specific (**Table1 and Supplementary Table 2**). Interestingly, while in WT MEFs we observed deletions with medium lengths of 5 to 9 nucleotides, shorter deletions of 1 to 4 nucleotides were found in FA-A MEFs (**Fig 2**). A more detailed analysis of the different indels generated at the on-target site (**EV2 and 3**) revealed subtle differences between samples of WT or FA-A MEFs, suggesting a potential different repair preference upon nuclease treatment in the two cell types. Furthermore, these analyses revealed that the on-target cleavage predominantly occurred in the range of nucleotides 222 to 226 (**EV4**).

**Fig 2.**
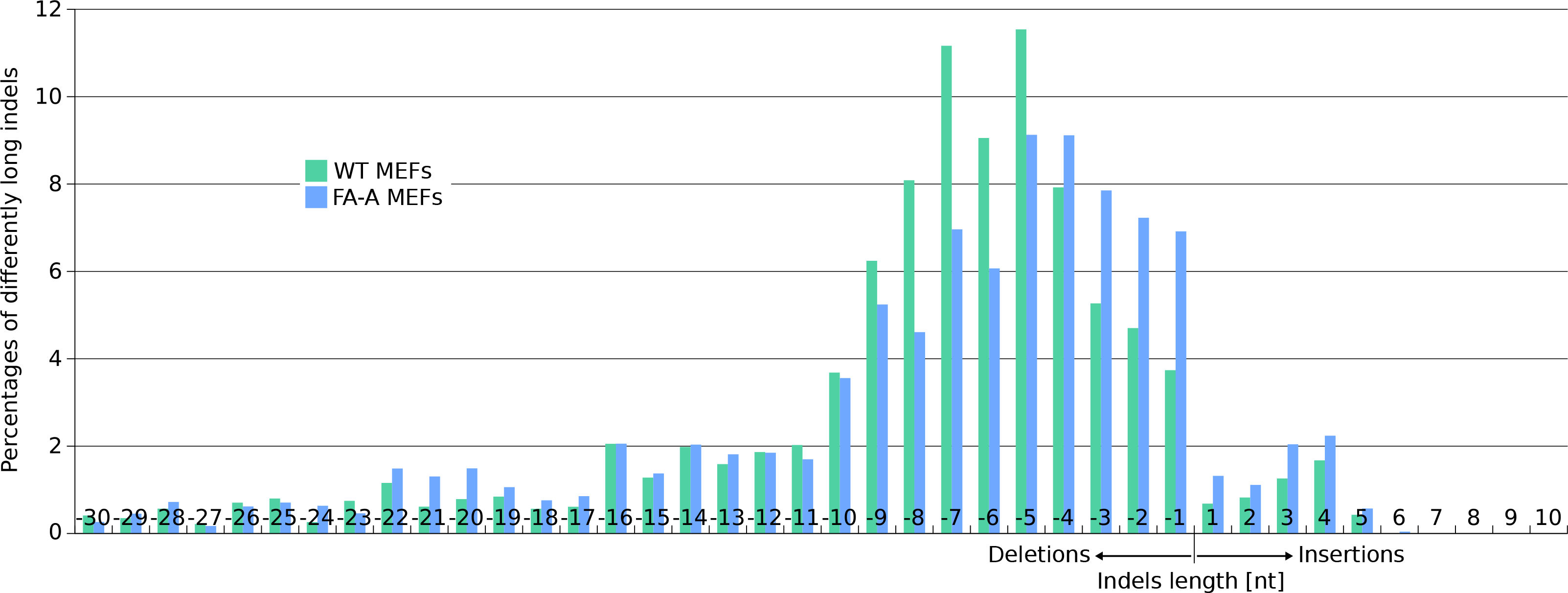
Differential length of nuclease-induced indels in WT vs. FA-A MEFs. All single deletions and insertions at the on-target site were analyzed for their length in nuclease treated WT- and FA-A MEF cells. The percentage of the different deletions or insertions was calculated in respect to all deletions or insertions respectively. The range shown was reduced to deletions with a maximal length of −30 nt and insertions with a maximal length of 10 nt.

### Generation of disease-free and functional FA-A MEF clones corrected by gene targeting in the *Mbs85* locus

Seamless integration of the donor into the *Mbs85* site was validated via Sanger sequencing of the PCR amplicon obtained from the 3’ integration junction analysis in one of the five selected clones (Clone #76#, **Fig 3A**). Furthermore, metaphase chromosomes were used to characterize the specific PGK-h*FANCA* transgene integration in chromosome 7 (where the *Mbs85* locus is located) by fluorescence *in situ* hybridization (FISH). We used one probe that recognized the two chromatids of chromosome 7 (which appear as two green spots pointed out by a green arrow in **Fig 3B**), and a second probe that recognized the PGK-h*FANCA* transgene (which appears as a red spot pointed out by a red arrow in **Fig 3B**). Notably, in the two clones analyzed (i.e. #7# and #76#), the red and green signals co-localized in the same metaphase spread, strongly suggesting that the targeted integration correctly occurred at the *Mbs85* locus (**Fig 3B**). However, non-targeted integration events were also observed in some of the metaphase spreads, such as in clone #76# (**Fig 3B**).

**Fig 3.**
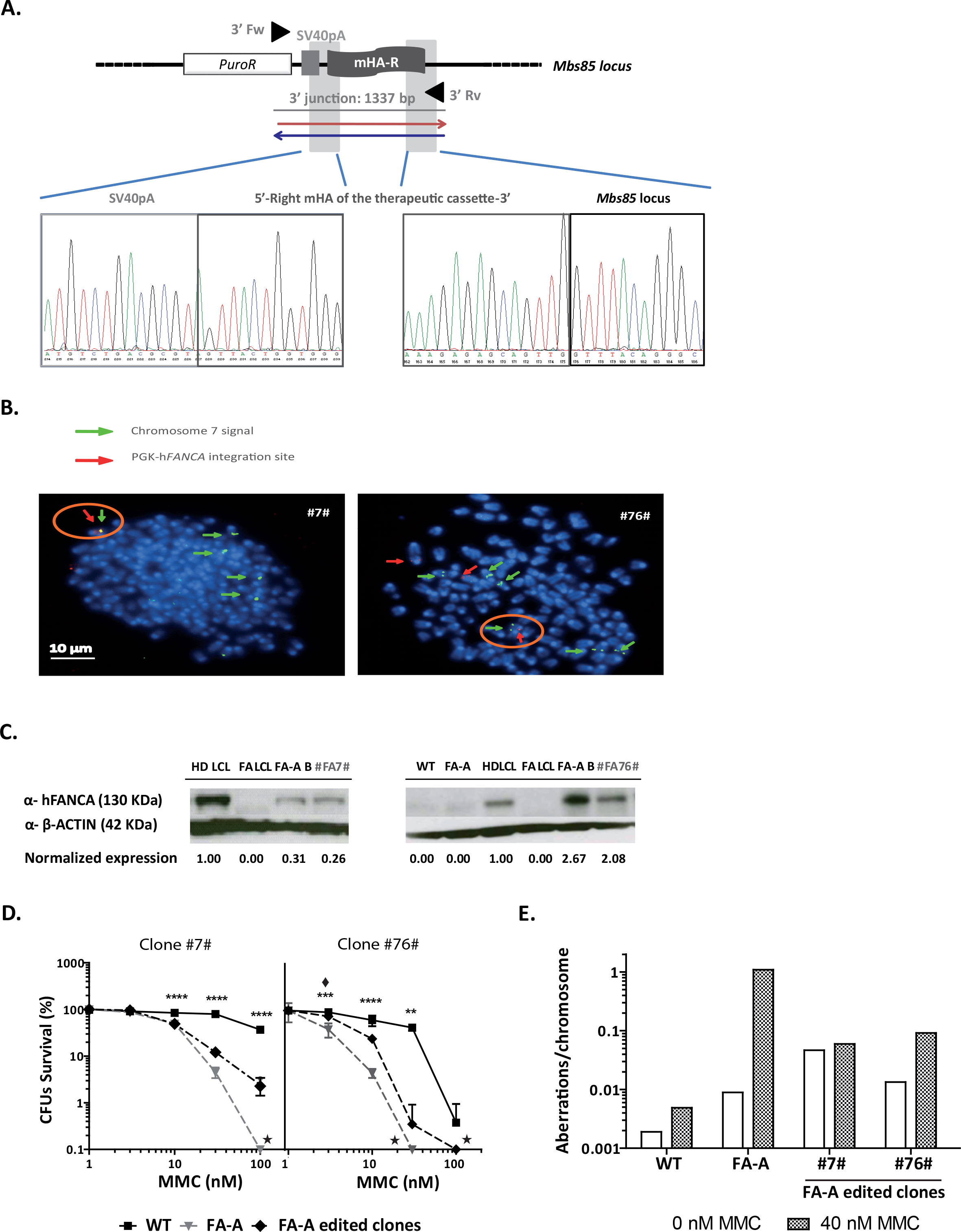
Targeted integration of the therapeutic h*FANCA* donor into the *Mbs85* locus and phenotypic correction of gene-edited FA-A MEF clones #7 and #76#. **A)** Sequenced region of clone #76# obtained with several primers displaying part of the chromatogram containing the SV40pA sequence, the left and right mHA of the therapeutic cassette, and part of the right location of the *Mbs85* locus (3’ integration junction). **B)** FISH analyses in gene-edited FA-A MEF indicating the integration of the therapeutic PGK-hFANCA donor (red spot signal and red arrows) into chromosome 7, (two green signals corresponding to the two chromatids indicated with a green arrow) where the *Mbs85* locus is located. Co-localization of both signals is indicated with an orange circle. DAPI chromosome staining is shown in blue. Scale bar represents 10 μm for all the microphotographs. From left to the right: targeted integration event in clone #7# and targeted and non-targeted integration events in clone #76#. **C)** Western blot analysis of hFANCA expression in the bulk of gene-edited FA-A MEFs, as well as in the gene-edited derived clones. Expression levels were calculated as a fold change with respect to β-ACTIN that was used as a loading control and then normalized against the hFANCA expression of lymphoblastic cells derived from a healthy human donor (HD LCL). FA LCL: lymphoblastic cells derived from a FA-A patient; FA-A: immortalized non-gene-edited FA-A MEFs; WT: immortalized non-gene-edited WT MEFs; FA-AB: bulk of edited FA-A MEFs (T2.5+D0.75). Analysed edited clones: #7# and #76#. **D)** Survival curves after exposure to different concentrations of MMC (0, 3, 10, 30, 100 nM) of edited FA-A MEF clones. In each curve the analysed clone is represented, together with untransfected WT and FA-A MEFs. Star (★) indicates that no colonies were generated in this condition at the corresponding doses of MMC. The survival shown would correspond to the growth of a single colony in these cultures. (**) P-value <0.01, (***) P-value <0.001, (****) P-value <0.0001 indicate significant differences with respect to FA-A group, with a twoway ANOVA followed by a post-hoc Bonferroni test. Asterisks indicate groups with differences (**\*** WT MEFs; ♦ Clone #76#) with respect to FA-A group. **E)** Chromosomal aberrations per chromosome induced by MMC, analysed in metaphases of gene-edited FA-A MEF clones, in comparison with non-edited WT and FA-A MEFs.

To assure the generation of functionally corrected gene-edited FA-A MEF clones we sought to verify whether these clones were functionally corrected in comparison with their parental non-edited FA-A MEFs. First, the functionality of the cassette was tested to determine the expression of hFANCA by Western blot in gene-edited clones. Since mouse MEFs do not express the human FANCA protein, we normalized the expression of hFANCA using a healthy human donor lymphoblastic cell line (HD LCL). As shown in **Fig 3C**, FA-A MEFs, WT MEFs and human FA-A LCLs did not show detectable levels of the hFANCA protein, as expected. However, we observed hFANCA expression both in the bulk population of gene-edited FA-A MEFs (2.5 µg of each TALEN monomer and 0.75 µg of donor) and in clones #7# and #76#. This result demonstrates that our gene-editing approach promoted the expression of the therapeutic h*FANCA* gene in FA-A MEFs upon integration of its expression cassette in the murine *safe harbor Mbs85* locus. As FA cells are characterized by their hypersensitivity to DNA interstrand cross-linking agents, such as mitomycin C (MMC), the sensitivity of uncorrected and gene-edited FA-A MEFs to this drug was tested. Cells were cultured with different doses of MMC and their survival was analyzed at 14 days post-treatment. Selected gene-edited clones #7# and #76# showed increased MMC resistance as compared to non-gene-edited FA-A MEFs (**Fig 3D**).

The generation of chromosomal breaks upon exposure to DNA cross-linking drugs is another characteristic of FA cells. Thus, to demonstrate the reversion of the characteristic phenotype of FA cells, chromosomal aberrations were studied in edited FA-A MEF clones were studies in the presence and the absence of MMC. Consistent with their restored FA pathway, gene-edited FA-A MEF clones presented a lower number of MMC-induced aberrations per chromosome in comparison with non-corrected FA-A MEFs. Interestingly, the number of chromosomal aberrations per MMC-treated cellwas 10.9-12.1 times lower in edited cells as compared to values obtained in their parental non-edited FA-A MEFs (**Fig 3E**). Altogether these results demonstrate that the specific integration of the therapeutic PGK-h*FANCA* cassette into the murine *safe harbor Mbs85* locus corrects the disease phenotype of FA-A MEFs.

### Efficient HDR-mediated gene editing in the mouse *Mbs85* locus of WT HPCs

Our laboratory has previously established an efficient and specific gene editing approach to correct fibroblasts and CD34^+^ cells from FA-A patients by harnessing the homology directed repair (HDR) pathway to integrate a therapeutic cassette into the human *AAVS1 safe harbor* locus of these cells (Diez et al, 2017; Rio et al, 2014). Experiments shown in Figures 1-3 have shown the feasibility of targeting the mouse ortholog of the human *AAVS1 gene* in mouse FA-A fibroblasts. In the subsequent experiments our goal was to test in mouse HSPCs the therapeutic relevance of the gene editing strategy described above to insert the human *FANCA* donor cassette into the murine *AAVS1* ortholog.

Lineage negative bone marrow (Lin^-^ BM) cells were purified by cell sorting and then pre-stimulated with a cocktail of hematopoietic cytokines (Navarro et al, 2006; Riviere et al, 2014) for 48 hours to promote cell cycling of quiescent HSPCs. Upon nucleofection with the TALEN and the therapeutic donor described above, pre-stimulated cells were maintained in culture in the presence of hematopoietic growth factors to boost HDR-mediated gene-editing repair (Branzei & Foiani, 2008) until performing the analyses at 48 hours post-nucleofection (**Fig 4A**).

**Fig 4.**
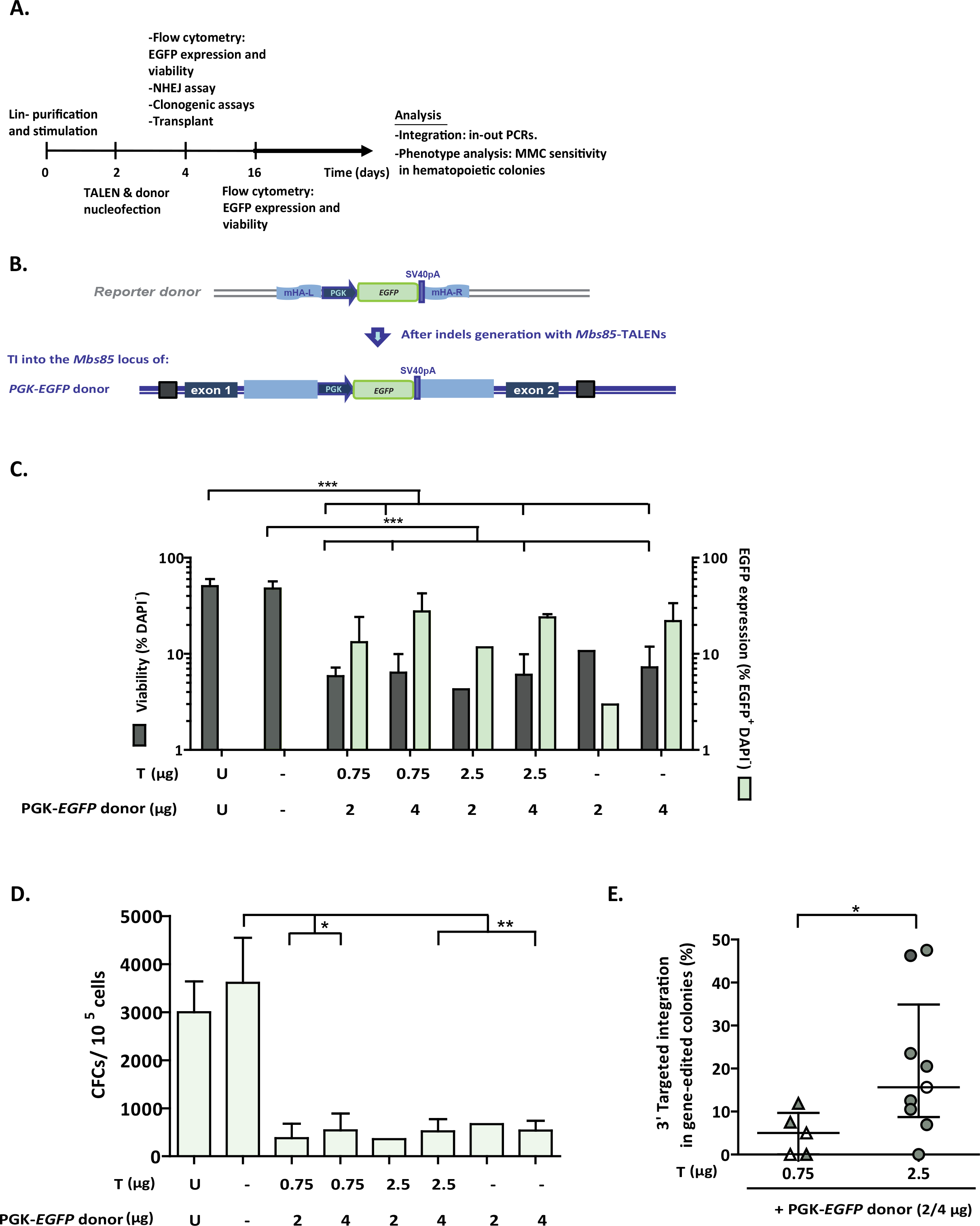
Gene-targeting in WT murine HSPCs. **A)** Flow chart indicating the study design and the different analyses performed in gene-edited murine HSPCs. **B)** Schematic showing the architecture of the murine *Mbs85* locus, with the target sites of the TALENs highlighted, and the structure of the donor used with the *EGFP* reporter cassette driven by the phosphoglycerate kinase promoter (PGK) flanked by sequences homologous to the genomic target locus. The resulting locus upon targeted integration (TI) of the donor is indicated in the lowest part of the panel. mHA-L and mHA-R: homology arms to the murine *Mbs85* locus; SV40pA: simian virus 40 polyA sequence; **C)** Analysis of viability (percentage of DAPI^-^ cells) and percentage of EGFP^+^ cells in WT Lin^-^ BM cells nucleofected with the TALEN and the PGK-*EGFP* reporter donor at 48 hours post-nucleofection. U: untransfected cells; Double negative (-): Nucleofected without DNA; T: different doses of TALEN monomers (from 0.75 to 2.5 µg); D: 2 or 4 µg of the PGK-*EGFP* reporter donor. Data are the mean ± S.D. (n=5 experiments). (***) P-value <0.001 indicates significant differences with a one-way ANOVA followed by a *post-hoc* Tukey test. **D)** Clonogenic assay to evaluate the ability of WT Lin^-^ BM cells to generate hematopoietic colonies under the conditions shown in C). Data are the mean ± S.D. (n=5 experiments). (*) P-value <0.05, (**) P-value <0.01 indicate significant differences with a one-way ANOVA followed by a *post-hoc* Tukey test. **E)** Targeted integration percentage of the PGK-*EGFP* reporter donor into the *Mbs85* locus of WT Lin^-^ BM cells for the 3’ integration junction calculated in the hematopoietic colonies that were positive for the PCR. Percentages calculated in nucleofected cells with different doses of TALEN (with triangles, 0.75 µg of each monomer; with circles, 2.5 µg of each monomer) and the donor (in white, the dose of 2 µg; in black, the dose of 4 µg). Data are the median ± interquartile range (n=5-9). (*) P-value <0.05 indicates significant differences with a Mann-Whitney test.

First, the activity of the *Mbs85*-specific TALENs was evaluated by the Surveyor assay in BM-derived Lin^-^ cells from WT and FA-A mice (**EV5A)**. The cleavage efficacies ranged from 30.4% ± 5.9% to 20.5% ± 10.1% respectively, indicating successful disruption of the target locus in HSPCs from these genotypes (**EV5B**). Since HSPCs are more sensitive to *in vitro* manipulation than adherent MEFs and because limited data are available on effective procedures to nucleofect murine HSPCs (Schiroli et al, 2017), we analysed the survival and transfection efficiency of mouse Lin^-^ WT BM cells 48 hours after nucleofection with different doses of TALEN plasmids (0.75 µg or 2.5 µg each monomer) and a PGK-*EGFP* reporter donor (2 µg or 4 µg, respectively), flanked by sequences homologous to the murine *Mbs85* locus (**Fig 4B**). A marked reduction (about 7-fold) in the viability of DNA nucleofected samples was observed with respect to untransfected or to mock transfected cells (**Fig 4C**).

When cells were nucleofected with different doses of TALEN and PGK-*EGFP* reporter donor, the percentage of EGFP^+^ cells at 48 hours post-nucleofection ranged from 11.7% to 27.6% (transient expression), regardless of the TALENs dose. In the absence of the TALENs, transient expression of EGFP in these cells ranged between 3% and 22% (**Fig 4C**). The ability of nucleofected WT Lin^-^ BM cells to generate hematopoietic colonies was also evaluated. DNA nucleofection reduced the ability of BM hematopoietic progenitors to generate colonies approximately 9-fold with respect to mock transfected cells (**Fig 4D**).

Although no EGFP-fluorescent colonies were observed in these cultures, we investigated the occurrence of specific integrations of the PGK-EGFP reporter donor into the *Mbs85* locus, by conducting PCRs for the 3’ integration junction in hematopoietic colonies. Strikingly, targeted integration (TI) frequencies of 4.8 ± 2.2% were measured in samples nucleofected with 0.75 µg of each TALEN monomer together with the PGK-EGFP donor, and this value increased to 20.9 ± 9.4% when the TALEN dose was increased to 2.5 µg (**Fig 4E**).

### Phenotypic correction of gene-edited FA-A HPCs

To prove the therapeutic potential of gene editing in FA-A HSPCs, the *EGFP* reporter donor used in WT cells was replaced with the therapeutic PGK-h*FANCA* donor used in the MEF experiments (**Fig 1 and 3**). Since higher TI frequencies were obtained in WT Lin^-^ cells using 2.5 µg of each TALEN monomer, we used this dose in subsequent experiments, together with 2 and 4 µg of the PGK-FANCA donor.

Nucleofection of FA-A Lin-BM cells with the TALEN and the donor plasmids reduced the viability of these cells 11-fold either when compared with non-nucleofected or mock nucleofected cells (**Fig 5A**). Similarly, DNA nucleofection also reduced the clonogenic ability of FA-A Lin^-^ BM cells approximately 9-fold in comparison with cells subjected to nucleofection in the absence of DNA (**Fig 5B**), which is similar to what we observed in nucleofected WT Lin^-^ BM cells (no significant differences were observed using a two-way ANOVA).

**Fig 5.**
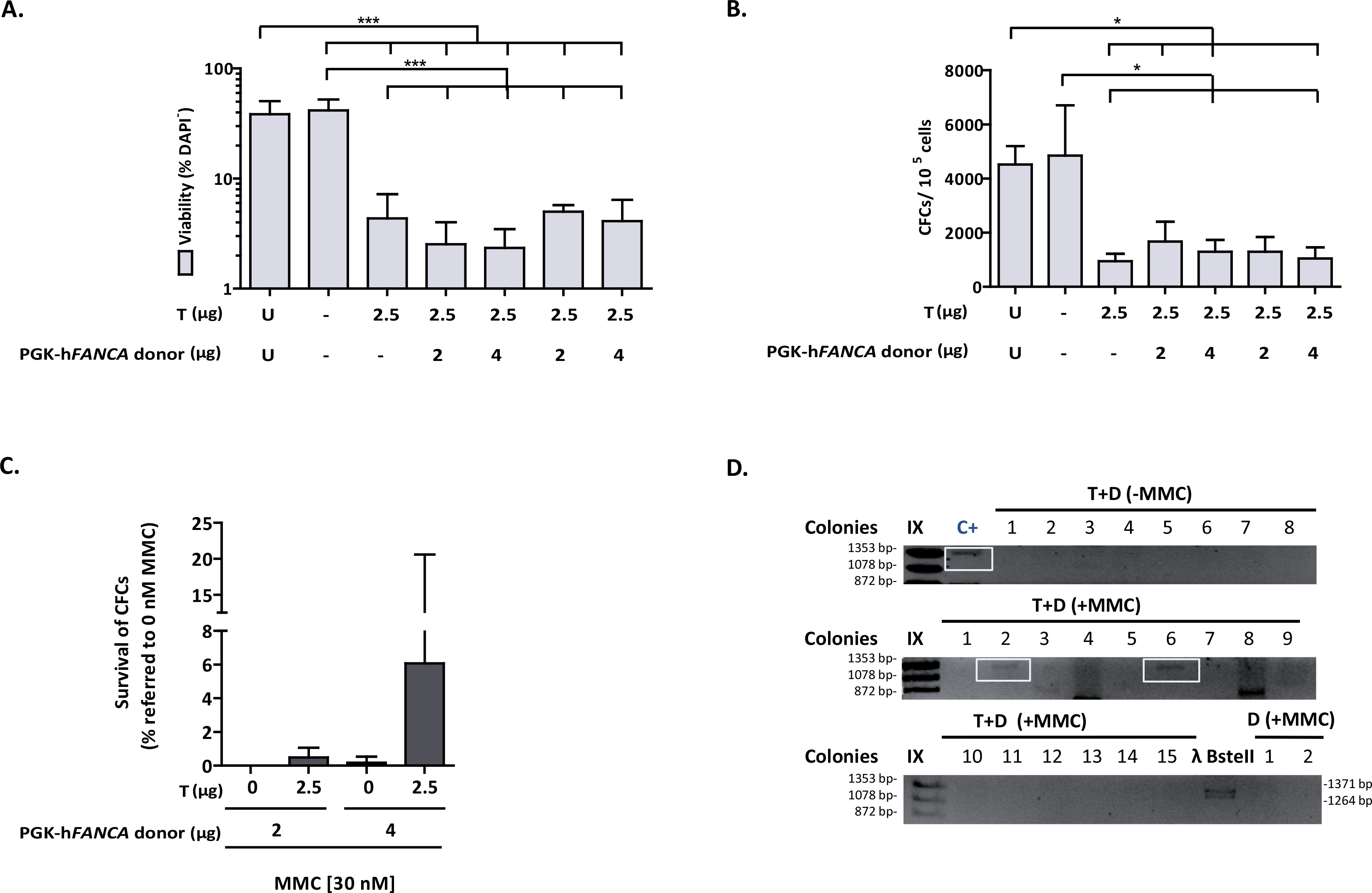
Gene-targeting in FA-A murine HSPCs. **A)** Analysis of viability (percentage of DAPI^-^ cells) in FA-A Lin^-^ BM cells nucleofected with the TALEN and the therapeutic PGK-h*FANCA* donor at 48 hours post-nucleofection. U: untransfected cells; Double negative (-): Nucleofected without DNA; T: 2.5 µg of each NN-TALEN monomer; D: 2 or 4 µg of the therapeutic PGK-h*FANCA* donor. Data are the mean ± S.D. (n=7 experiments). (***) P-value <0.001 indicates significant differences with a one-way ANOVA followed by a *post-hoc* Tukey test. **B)** Clonogenic assays of nucleofected Lin^-^ cells under the conditions shown in A). Data are the mean ± S.D. (n=7 experiments). (*) P-value <0.05 indicate significant differences with a oneway ANOVA followed by a *post-hoc* Tukey test. **C)** Phenotypic correction measured with clonogenic assays in nucleofected FA-A Lin^-^ BM cells treated with 30 nM MMC compared to cells cultured in the absence of MMC. MMC survivals are indicated considering the number of hematopoietic colonies generated without drug selection as 100%. Cells were subjected to nucleofection with 2.5 µg of each TALEN monomer and the therapeutic PGK-h*FANCA* donor (2 and 4 µg). Data are presented as mean ± S.D. (n=4 with the except of D2 µg where n=1 experiments). No statistical differences were found among groups with a non-parametric Kruskal-Wallis and median test. **D)** Representative PCR analysis for the study of the 3’ integration junction (1,337 bp). C+: sequenced genomic DNA positive for the integration junction in edited FA-A MEFs; T+D: samples from nucleofected FA Lin^-^ BM cells with 2.5 μg of each TALEN monomer together with 4 μg of the therapeutic PGK-h*FANCA* donor; D: 4 μg of the therapeutic PGK-h*FANCA* donor; IX and λ BstII: DNA molecular weight markers. Analysed colonies are numbered and the positive ones framed in blue.

Remarkably, when cells were plated in methylcellulose in the presence or absence of 30 nM MMC, 6% of the hematopoietic colonies corresponding to samples nucleofected with the TALEN and the therapeutic donor survived in the presence of MMC, while only 0.2% of colonies survived in MMC when only the donor was used, indicating that targeting the integration of the therapeutic transgene to *Mbs85* corrected the hypersensitivity of FA HSPCs to MMC (**Fig 5C**). In 16.3% of the colonies that grew in the presence of MMC we confirmed the amplification of the expected Mbs1 3´ integration junction (**Fig 5D**), thereby indicating that correction of MMC-hypersensitivity in primary FA-A mHSPCs was a consequence of the PGKh*FANCA* targeted integration in the *Mbs85* locus.

## DISCUSSION

The *AAVS1* locus has been defined as a bona-fide s*afe harbor* locus in humans (Lombardo et al, 2011), in which an exogenous gene could be efficiently expressed in various cell lines and iPSCs (DeKelver et al, 2010; Dreyer et al, 2015; Hockemeyer et al, 2009; Li et al, 2017; Lombardo et al, 2011; Mizutani et al, 2016; Mizutani et al, 2015; Oceguera-Yanez et al; Ordovas et al, 2015; Ramachandra et al, 2011; Smith et al, 2008; Zou et al, 2011). Therefore, in this work, we have decided to carry out a targeted integration strategy in the mouse *Mbs85* orthologous locus in order to asses if the corresponding mouse locus may serve as a new site where gene editing can be performed in mouse disease models such as Fanconi anemia (FA).

Gene editing of HSPCs is an attractive strategy in FA due to the *in vivo* proliferation advantage of corrected HSPCs over non-corrected ones (Gregory et al, 2001; Gross et al, 2002; Lo Ten Foe et al, 1997; Mankad et al, 2006; Rio et al, 2017; Soulier et al, 2005; Waisfisz et al, 1999). TALEN delivery has remained one of the major obstacles in providing short-term and dose-controllable nuclease activity based on previous studies (Cai et al, 2014; Holkers et al, 2013; Liu et al, 2014; Mock et al, 2014). We nucleofected a pair of TALENs in combination with a homologous donor template, in the form of plasmid DNA, in both MEFs and HSPCs from wt or FA-A mice. Carrying out targeted integration in FA-A cells constitute challenging approaches of gene therapy due to the involvement of the FA proteins in promoting HDR (Adamo et al, 2010; Nakanishi et al, 2005), which could affect the efficiency of the homologous integration of the therapeutic cassette in the targeting site of the genome.

Once the expression of TALEN monomers in HEK293T cells was confirmed (**EV1B**), we established proof of principle for gene editing in FA-A MEFs. Using a stepwise experimental approach as followed in previous gene-targeting studies (Bednarski et al, 2016; Diez et al, 2017; Rahman et al, 2015), we observed relatively high transfection efficiencies (mean value of 30%) and viabilities (ranging from 47% to 78%), indicating that in this cell type cytotoxicity was not a limiting factor to perform gene editing. We also observed that when a donor template was used together with the TALENs, the percentage of repair by NHEJ was significantly reduced; suggesting that a proportion of NHEJ repair was replaced by HDR in the presence of a donor template.

A thorough analysis of the off-target effects is crucial for the further development of gene-editing based therapeutics. We used high-throughput sequencing to assess the off-target sites previously predicted *in silico* with the PROGNOS software (Fine et al, 2013) in nucleofected MEFs, as this method is the preferred one to detect indel mutations induced at low frequencies with great sensitivity (Hendel et al, 2015; Koo et al, 2015). Indels generated by the nucleases were highly specific in WT and FA-A MEFs, as demonstrated by the high percentage of on-target indels and the low percentage of off-target generated indels (**Table 1**). As previously shown for other TALEN (Mussolino et al, 2014; Mussolino et al, 2011), our study confirms the high specificity of the *Mbs85*-specific TALENs.

**Table 1.**
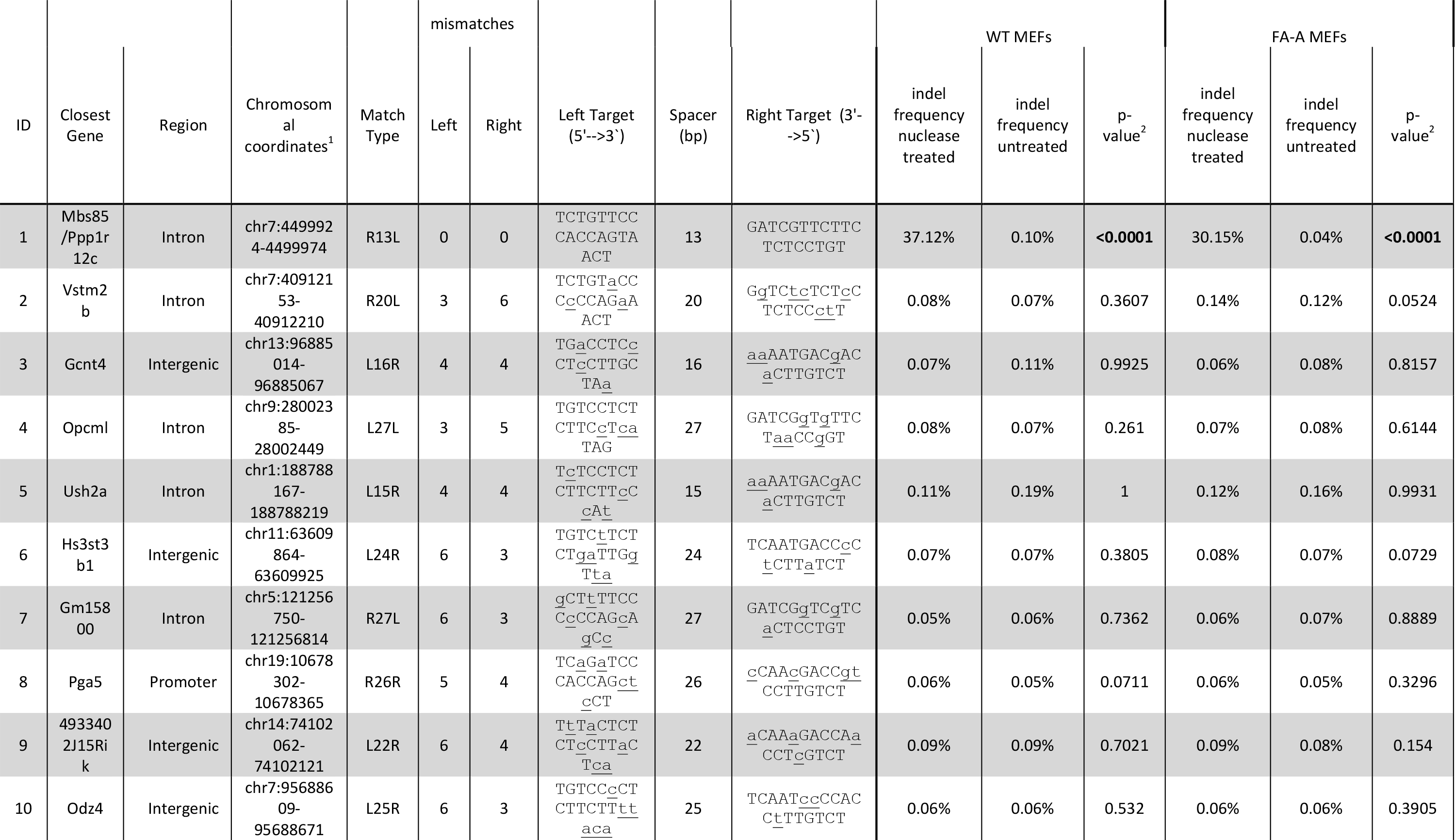
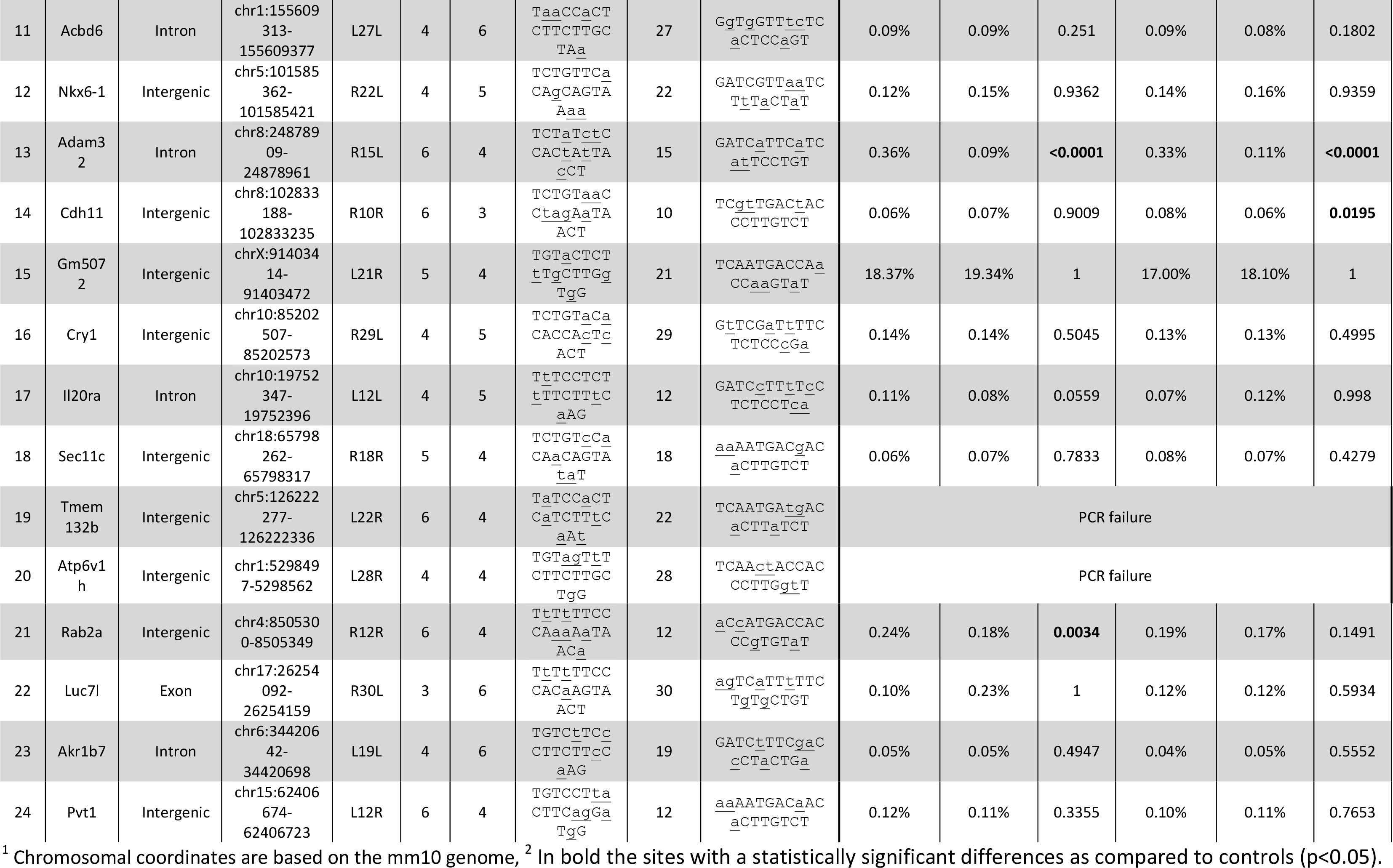
Off-target cleavage of Mbs85-specific TALENs. The top off-targets are listed according to the ranking generated with TALENv.2 algorithm from PROGNOS software. The nucleotide differences in the left and in the right targets, the closest gene, the region and chromosomal coordinates of the off-target location are indicated. The match type and the sequence of the left and the right target are also included. The indel frequency and p-value are also indicated.

In some colonies, we had difficulties amplifying the specific PCR bands corresponding for both integration junctions. This could be due to the efficiency of the PCRs, or to the limited and poor quality of the DNA obtained from single colonies. However, as has been previously mentioned, other mechanisms of repair of the DSBs could have been involved apart from HRR, giving rise to different outcomes at the different junctions. Importantly, most of the colonies that arose from samples only nucleofected with the donor did not amplify any of the specific integration bands. In the few cases in which this was observed, on-target non-directed integration events independent of TALEN cleavage may have occurred. We confirmed specific integration of the therapeutic PGK-h*FANCA* cassette into the *Mbs85* locus by PCR when both the donor and the TALENs were nucleofected simultaneously, indicating the generation of HDR-mediated integrations of the therapeutic donor at the on-target site (**Fig 1E**). Importantly, the specific integration was confirmed by Sanger sequencing (**Fig 3A**) and FISH (**Fig 3B**). However, we also observed the occurrence of non-targeted integration events, highlighting the risks associated with DNA delivery (**Fig 3B**, clone #76#).

In this study, we also confirmed the efficient phenotypic correction of edited MEF cells. This conclusion was deduced both from the functional expression of the hFANCA protein (**Fig 3C**) and the MMC resistance (**Fig 3E**) and reduced MMC-induced chromosomal instability (**Fig 3F**) of edited clones. Taken together, these studies demonstrate the correction of the FA/BRCA pathway through TALEN-mediated targeted integration of a therapeutic h*FANCA* cassette into the *Mbs85* locus of FA-A MEFs.

Once we proved the efficacy of our gene-targeting experiments in FA-A MEFs, we performed similar experiments in mouse HSPCs in order to demonstrate the feasibility of a gene-targeting strategy in the *Mbs85* locus of these cells using the PGK-*EGFP* reporter donor. Working with sorted Lin^-^ BM cells from WT mice, we observed the expression of the integrated *EGFP* (**EV6**), facilitated by the targeted integration of the *EGFP* cassette into the *Mbs85* locus. Despite the absence of EGFP expression in *Mbs85-*edited WT HSPCs, targeted integration in up to 20.98 ± 9.4% of clones were observed using 2.5 µg of each TALEN monomer together with 4 µg of the *EGFP* reporter donor (**Fig 4E**). This observation and the very low proportion of EGFP-expressing cells observed in liquid cultures suggests a restricted expression of transgenes in the *Mbs85* locus of mHSPCs as has previously been reported in the human *AAVS1* locus of hESCs (Ordovas et al, 2015).

Consistent with the results observed in WT Lin^-^ BM cells, we demonstrated for the first time the feasibility of conducting a therapeutic gene targeting approach into the *Mbs85* locus of primary Lin^-^ BM cells from FA-A mice using the PGK-h*FANCA* donor. As for WT mHSCs, both the viability and clonogenic potential of nucleofected FA-A mHSCs (**Fig 5A** and **B**) declined sharply. Regarding the targeted integration observed in WT as compared to FA HSPCs, similar efficacies were observed, supporting the hypothesis that although HDR could be moderately affected in FA-A cells (as was observed in a recent study from our laboratory (Diez et al, 2017)), gene editing is feasible in these cells, due to their mild HDR defects, in comparison with FA-D1 cells.

Of particular importance is the fact that our results show the correction of the MMC-hypersensitivity phenotype in primary FA-A mHPCs. Correctly edited FA-A HPCs survived cytotoxic concentrations of MMC, similarly to what was observed in FA-A MEFs, highlighting the therapeutic potential of the proposed gene therapy approach. Although *Mbs85* might limit the efficacy of expression of integrated cassettes in this locus, the achievement of significant levels of MMC-resistant in edited FA-A CFCs is consistent with our previous observations showing that low levels of FANCA can result in a therapeutic effect (Almarza et al, 2007; Gonzalez-Murillo et al, 2010).

In our studies of DNA nucleofection in mouse HSPCs, we observed a marked cytotoxicity due to transferring plasmid DNA to these cells, as mock nucleofection showed no impairment in viability (**Fig 4D and 4E**). This toxicity that results in a reduction in the efficiency of these strategies, currently constitutes one of the main limitations to conduct *ex vivo* therapeutic approaches of gene editing in hematopoietic diseases (Cornu et al, 2017), however, other delivery methods might be a safer and overcome this issue (De Ravin et al, 2016; Diez et al, 2017; Mock et al, 2015; Poirot et al, 2015; Qasim et al, 2017; Wang et al, 2015). Nevertheless, our results showing toxicity of gene editing approaches in mouse HPCs are consistent with previous studies aiming at the gene targeting in primary mHPCs (Gundry et al, 2016; Riviere et al, 2014; Schiroli et al, 2017) and confirm that gene editing in the murine HSPC compartment is challenging, particularly in cells capable of hematopoietic repopulation.

Overall, our data provide evidence of successful therapeutic gene editing in the mouse *Mbs85* orthologous locus in fibroblasts and HPCs of a mouse FA model, and establishes the rationale and the proof of principle to use this locus for gene correction in other diseases in preclinical studies to investigate the efficacy and safety of future gene editing strategies.

## MATERIALS AND METHODS

### Cells and cell culture

HEK-293T (ATCC-CRL-3216™) cells were cultured in DMEM 1X with GlutaMAX^TM^ (Gibco) with 10% Hyclone (GE Healthcare) and 1% Penicillin/Streptomycin (P/S) (Gibco).

MEFs both from FVB/NJ WT and FVB FA-A mice were obtained from the chorion of 13.5 E pregnant females as has been described previously (Navarro et al, 2014). MEFs from WT or FAA mice were immortalized by a transient transfection with pLXSN 16 *E6E7* and pCL-ECO-gag-pol (Naviaux et al, 1996) using the CaCl_2_ DNA precipitation method and cultured also in DMEM 1X with GlutaMAX^TM^ (Gibco) with 10% Hyclone (GE Healthcare) and 1% Penicillin/Streptomycin (P/S) (Gibco).

BM cells were isolated from FVB/NJ WT, FVB FA-A or C57BL/6J mice by flushing the femurs and tibias of these mice in IMDM (Gibco). The cellular suspension was incubated with lysis solution (CINH_4_ with CO_3_HK 1M with EDTA 0.5M) (Merck KGaA) for 5 minutes at RT in darkness. Then, cells were washed with PBS 1X (Sigma^®^ Life Sciences) with 5% Hyclone and 5% P/S. Purified mHPCs, Lin^-^ (lineage-negative) cells, were obtained by whole BM cell sorting by immunoselection. Lin^-^ cells were expanded in StemSpan™ (StemCell™ Technologies) with 1% GlutaMAX™ with growth factors and cytokines and 1% P/S. The following factors were added: 100 ng/ml mouse stem cell factor (mSCF), 100 ng/ml human interleukin 11 (hIL-11), 100 ng/ml human FMS-like tyrosine kinase 3 ligand (hFlt3), 100 ng/ml human thrombopoietin (hTPO)(EuroBioSciences).

With the exception of HEK-293T cells that were cultured at standard normoxic (21% O_2_-5% CO_2_) conditions the rest of the cells were cultured in hypoxia (5% O_2_-5% CO_2_) at 37ºC, and 95% relative humidity.

### TALENs, donors and control plasmids

TALE-based DNA binding domains were assembled using Golden Gate assembly kit (Morbitzer et al, 2011) modified based on our previously optimized TALEN scaffold (Mussolino et al, 2011) (Δ135/+17). The vectors used (Mussolino et al, 2014) included the 17.5th repeat and the wild-type *FokI* cleavage domain (pVAX_CMV_TALshuttle(xx); ‘xx’ stands for the four different 17.5th RVDs used, NI, NG, HD and NN).

Donor plasmids were flanked by two homology arms (HA) of the *Mbs85* locus of 806 bp and 860 bp, respectively. The therapeutic donor contained a PGK.FANCA.E2A.PuroR.SV40pA fragment that was chemically synthesized by GenScript. PGK-*EGFP* reporter donor was cloned in the backbone of the therapeutic cassette by digestion of the PGK.*EGFP* fragment from the pCCL.PGK.*EGFP*.wPRE* plasmid with EcoRI/NotI restriction enzymes (New England Biolabs, Ipswich, Massachusetts, USA). A PGK-h*FANCA* donor with longer homology arms, was used as a positive control for targeted integration in the *Mbs85* locus

### Lipofection and nucleofection

HEK-293T cells were lipofected at 70% confluence one day after seeding 1×10^5^ cells with Lipofectamine^®^ 2000 Reagent according to manufacturer’s protocol (Invitrogen). 400 ng of DNA of each TALEN monomer were co-transfected with 100 ng of an *EGFP* control plasmid and 500 ng of pUC118 control plasmid.

For nucleofection of WT or FA-A immortalized MEFs, 2×10^6^ cells per condition were used with the Amaxa MEF2 Nucleofector^®^ Kit (Lonza Group) using program T20 of the Nucleofector™ I device, 2.5 µg of each TALEN monomer were nucleofected together with different donor doses (0.75, 2 and 4 µg). Enrichment during 5 passages with puromycin (1-1.25 µg/ml) was performed, then clones were generated by limiting dilution to perform gene targeting studies

For the BM hematopoietic cells, 1.4×10^6^ cells of purified BM Lin^-^ cells were nucleofected with an *EGFP* control plasmid or the corresponding doses of the TALEN monomers and donors after pre-stimulation during 48h. Lin^-^ BM cells were nucleofected with the 4-D Nucleofector™ device using the P3 Primary Cell 4D-Nucleofector^®^ X Kit (Lonza Group) kit with the ED-113 program.

### Cell sorting & flow cytometry

Lin^-^ cells were purified from C57BL/6J and FA-A FVB/NJ mice by cell sorting using lineage-specific antibodies phycoerytrin conjugated (BD Pharmingen) (anti-B220 (CD45R), anti-Mac-1 (CD11b), anti-Gr1 (Ly6G/C), anti-CD3-ε, and anti-Tert-119 antibodies) at a concentration of 0.06 μg/mL and 0.02 μg/mL, respectively. For the identification of LSK cells (Lineage negative, Sca-1^+^, c-Kit^+^), Lin^-^ cells were stained with the described cocktail, and with 0.06 μg/mL of anti-Sca-1-APC-Cy7 (BioLegend) and 0.15 μg/mL of anti-c-Kit-A647 antibodies (Southern). 4’,6-Diamidino-2-phenylindole (DAPI; Roche)-negative staining was used as a marker of cell viability. Analyses were performed in the LSR Fortessa cell analyser (BD/Becton, Dickinson and Company). Transfection efficiency was also determined by FACS analysis 48h post nucleofection. Off-line analyses were conducted with the FlowJo Software v7.6.5 (© FlowJo, LLC).

### Clonogenic -colony forming cell (CFC)- assays of hematopoietic progenitors and colony forming units (CFUs)

CFC assays were performed following manufacturer´s recommendations (Stem Cell Technologies, Vancouver, Canada). MMC (at 10 and 30 nM) (Sigma^®^ Life Sciences) and puromycin (1 µg/ml) (Sigma^®^ Life Sciences) was added, respectively, to analyze the number of gene-edited cells. 2×10^5^ cells were seeded at 48 hours post-nucleofection. Colonies were scored in an inverted microscope (Olympus IX70 WH10X/22). Cultures were maintained in hypoxia at 37°C, and 95% relative humidity.

Survival of immortalized and targeted FA-MEFs was analyzed by scoring the number of colonies derived from 200 cells (CFUs, colony forming units) exposed to increasing concentrations of MMC (0, 3, 10, 30, 100 and 300 nM). After one week, the medium was changed with fresh medium containing the same concentration of MMC. Cell viability was determined 14 days after cell seeding. All cultures were maintained in hypoxia at 37°C, and 95% relative humidity.

### *Cel I* (Surveyor) analysis of TALENs pairs activity

DNA was extracted using NucleoSpin^®^ Tissue kit (Macherey-Nagel) from cellular pellets obtained at 48h or after cell expansion. PCR fragments spanning the *Mbs85* TALENs target site were generated with primers mAAVS1 CelIF and mAAVS1 CelIR (**Supplementary Table3**). PCR was performed as follows: 200 ng of gDNA, 1.25 µl 10 µM of each primer, 0.5 µl 100 nM dNTPs, 10 µl of the Buffer 10X and 1 µl of Herculase enzyme (Herculase II Fusion Enzyme, Agilent Technologies) in a final volume of 50 µl. Cycling conditions were the following: 95°C for 2 minutes, 40 cycles of 95°C for 20 seconds, 56°C for 20 seconds and 72°C for 30 seconds, and finally one cycle of 72°C for 3 minutes. PCR products were purified with the NucleoSpin^®^ Gel and PCR Clean-up kit (Macherey Nagel) and amplified following the manufacturer’s instructions. Final products were migrated and revealed according to manufacturer instructions and analyzed in a Molecular Imager^®^ GelDoc™ XR+ System (BioRad). Quantification of the bands was performed using the Quantity One Software (BioRad).

### On-target integration analyses

Genomic DNA from either the bulk cell population or from FA-A MEF clones was extracted using NucleoSpin^®^ Tissue kit. gDNA from single colonies derived from CFCs assays was extracted as previously described (Charrier et al, 2011). Two PCR reactions were conducted both for the 5’ and the 3’ integration junctions of the *Mbs85* integration site. To analyze the integration of the PGK-h*FANCA* donor the following primers were used: the pair mAAVS1-5’F (1 & 2) and mAAVS1-5’R (1 & 2); and the pair mAAVS1-3’F (1 & 2) and mAAVS1-3’R (1 & 2) (**Supplementary Table3**). To analyze the integration of the PGK*-EGFP* donor: the pair mAAVS1-5’F3 and mAAVS1-5’R3, and the pair mAAVS1-EGFP-3’F and mAAVS1-EGFP-3’R were used. PCR reactions were conducted using 200 ng of gDNA from bulk cell population or single MEF clones, or 15 µl of gDNA from single colonies. PCRs from the 5’ and the 3’ integration junctions were performed with 1.25 µl at 10 µM of each primer, 0.5 µl at 100 nM dNTPs, 10 µl of the Buffer 10X and 1 µl of Herculase enzyme in a final volume of 50 µl. Cycling conditions were the following: 95°C for 10 minutes, 40 cycles of 95°C for 30 seconds, 59<u>°</u>C -mAAVS1-5’F1&R1-; or 58°C -mAAVS1-5’F2&R2-; or 62°C -mAAVS1-5’F3&R3- for 60 seconds (for the 5’ integration junction depending on the primers used); or 59°C -mAAVS1-3’F1&R1-; 61°C -mAAVS1-3’F2&R2-; 62°C -mAAVS1-EGFP-3’F&R- for 60 seconds (for the 3’ integration junction depending on the primers used); and 72°C for 1.5 minutes; and finally one cycle of 72°C for 10 minutes in a final volume of 50 µl.

PCR products of the 3’ integration junction performed in the FA-A MEF bulk population were sequenced by Sanger method. Primers used for sequencing were the followings: mAAVS1-3’-F2, mAAVS1-3’-R2; SeqAAVS1-1_3’_F; SeqAAVS1-2_3’_F; SeqAAVS1-3_3’_F; SeqAAVS1-4_3’_F; SeqAAVS1-1_3’_R (**Supplementary Table3**). Analyses of the sequences were done with Finch TV version 1.4.0 and Vector NTI software.

### Fluorescence in situ hybridization (FISH)

Cells were first arrested in metaphase with KaryoMAX Colcemid solution (Gibco) and then harvested after a treatment with a hypotonic salt solution. Two sets of probes were used to localize plasmid integration site. RP23-336p21 and RP24-129K5 specific bacterial artificial chromosomes (BACs) that map to the E3 band of the mouse chromosome 7 (Human BAC Clone Library, Children’s Hospital Oakland Research Institute [CHORI]) were used as controls. PGK-h*FANCA* transgene integration site was detected using as probe the DNA from the plasmid vector. Following the manufacturer’s specifications, BACs DNA was directly labelled by nick translation (Vysis) with SpectrumGreen-dUTPs, whereas plasmid DNA was labelled with SpectrumOrange-dUTP (Vysis). The probes were blocked with Cot-1 DNA and DNA sheared salmon sperm (Vysis) to suppress repetitive sequences, and hybridized overnight as 37°C onto metaphase spreads. After post-hybridization washes, the chromosomes were counterstained with DAPI in Vectashield mounting medium (Vector Laboratories). Cells images were captured using a cooled charge-coupled device (CCD) camera (Photometrics SenSys camera) connected to a computer running a Chromofluor image analysis system (CytoVision, Leica Biosystems).

### Western Blot

Western blot (WB) analyses were performed to determine the expression of the HA epitope of TALENs in HEK-293T cells using anti-HA antibody (NB600-363, Novus Biologicals). Human FANCA protein was also analyzed in immortalized FA-A MEFs and in gene-edited FA-A MEFs as previously described (Raya et al, 2009; Rio et al, 2014).

### Chromosomal instability assay

Gene-edited FA-A MEFs were subjected to medium containing 40 nM of MMC for 24 hours to induce DNA damage. Then, 0.05 µg/ml of colcemid (KaryoMAX Colcemid Solution, Gibco) was added to the cells followed by treatment with 0.56 % KCl during 15 min at 37°C and fixed in methanol: acetic acid (3:1). Extensions were made at 25°C with 48% humidity in a Thermotron chamber. Metaphases were stained with 10% Giemsa stain, giemsa’s azur eosin methylene blue solution (Merck) with Gibco^®^ Gurr Buffer Tablets (Gibco). Then, cells were placed in glass coverslips with a pair of droplets of Entellan (Merck). In general, a minimum of 20 metaphases per culture were studied for the analysis of aberrations. Cell images were taken with an inverted microscope (Olympus CK30).

### Off-target analysis

Possible off-target sites for the TALENs pair targeting the sequences 5’-TGTCCTCTCTTCTTGCTAG and 5’-AGTTACTGGTGGGAACAGA within the mm10 genome were predicted using the online tool PROGNOS (Fine et al, 2013; Lin et al, 2014). In the submitted query, 6 mismatches for each TALEN-half-site and a spacing distance between 10 bp and 30 bp were considered, together with hetero- and homodimeric sites (indicated in **Supplementary Table1**). The 24 top ranked off-targets (TALENv2.0 algorithm from PROGNOS) were analyzed by deep sequencing.

WT and FA-A MEFs were either treated with corresponding nucleases or left untreated. Genomic DNA was isolated to PCR amplify all predicted off-target loci (50 – 100 ng DNA were used per target; primers are listed in **Supplementary Table1**). Libraries were prepared from PCR Amplicons using the NEBNext^®^ Ultra™ II DNA Library Prep Kit for Illumina^®^ (NEB, E7645) and quantified using the ddPCR™ Library Quantification Kit for Illumina TruSeq (biorad, #186-3040). All samples were sequenced on an Illumina MiSeq platform with a MiSeq Reagent Kit v2, 500cycles (Illumina, MS-102-2003).

Paired-end reads from MiSeq reactions were quality trimmed by an average Phred quality (Qscore) greater than 20 using TrimGalor (www.bioinformatics.babraham.ac.uk/projects/trim_galore) and merged into a longer single read with a minimum overlap of 30 nucleotides using Fast Length Adjustment of SHort reads (FLASH)(Magoc & Salzberg, 2011). Merged sequences were aligned against all reference sequences (**Expanded View Table 1**) using Burrows-Wheeler Aligner (BWA). Alignments were analyzed for insertions and deletions within a range of ±20 bp of the predicted nuclease cleavage site.

### Statistical analyses

Statistical analyses were performed using GraphPad Prism version 5.0 for Windows (GraphPad Software). Results are shown as the mean ± Standard Desviation (SD), mean ± Standard Error of the Mean (S.E.M) in the case of the specific site integration in hematopoietic progenitors, and as the median ± interquartile range, from at least 3 replicates and from different experiments. When two sets of data were compared, two-tailed Student’s t-test or Mann-Whitney tests were performed depending on whether or not the values followed a normal distribution. Statistical significance of the indel frequency occurred at predicted off-target sites was determined using a one-tailed, homoscedastic Student’s t-test. This test was performed to compare the values that were above those obtained in untreated cells. When more than two sets of data were compared, a parametric one-way ANOVA followed by post-hoc multiple comparison Tukey test or a non-parametric Kruskal-Wallis followed by a post-hoc Dunn’s multiple comparison test were used. In some experiments, more than one variable was analyzed. In these cases, when the data suited a normal distribution, the two-way ANOVA test was applied followed by a Bonferroni post-hoc test. Significant differences were indicated as P-value <0.05 (*), P-value <0.01 (**), P-value <0.001 (***) and P-value <0.0001 (****).

## ACKNOWLEDGMENTS

This work was supported by grants from the “7th Framework Program European Commission (HEALTH-F5-2012-305421; EUROFANCOLEN)”, “Ministerio de Sanidad, Servicios Sociales e Igualdad” (EC11/060 and EC11/550), “Ministerio de Economía, Comercio y Competitividad y Fondo Europeo de Desarrollo Regional (FEDER)” (SAF2015-68073-R), “Fondo de Investigaciones Sanitarias, Instituto de Salud Carlos III” (RD12/0019/0023) and the German Federal Ministry of Education and Research (BMBF-01EO0803). The authors would like to thank Miguel A. Martin for the careful maintenance of NSG mice and Omaira Alberquilla for her technical assistance in flow cytometry. The authors also thank the Fundación Botín for promoting translational research at the Hematopoietic Innovative Therapies Division of the CIEMAT. CIBERER is an initiative of the “Instituto de Salud Carlos III” and “Fondo Europeo de Desarrollo Regional (FEDER)”. Finally, although this work was entirely performed with mouse cells, the authors feel deeply grateful to the FA patients and families, clinicians and to Aurora de la Cal as the secretary of the Spanish FA network for always instilling motivation and providing an example for work in this disease.

## AUTHOR´S CONTRIBUTION

MJ PB, SN, JB and CM: Conceived and designed the experiments. MJ PB, SN, YG, MV, MH, RSD, SRP, RP: Collection and assembly of data. CM, JS, PR, TC, JB: Provision of reagents, materials, analysis tool, ideas. MJ PB, SN, JB and CM: Manuscript writing.

## THE PAPER EXPLAINED

### Aim

Despite the success of GT trials that are at present currently undergoing with integrative vectors, gene-targeting approaches represent the next future and are proposed as safer alternatives. These strategies are based on the use of artificial nucleases that generate double strand breaks at specific locations in the genome which can be repaired by the homologous recombination pathway of the cells. Specific sites of the cell genome such as safe harbor locus are being targeted to exploit this approach and facilitate the integration of donor templates that can be therapeutic in the case of genetic diseases.

In this study we aimed at the insertion of reporter and therapeutic donors into the mouse *Mbs85* locus, ortholog of the human *AAVS1 safe harbor* locus using TALEN to investigate the feasibility of targeting a safe harbor locus in the murine context. This strategy was implemented in the *Fanca*^-/-^ (FA-A) mouse model for Fanconi anemia.

### Results

We have first demonstrated the activity of designed TALEN to cleave in the *Mbs85* locus of both FA-A embryonic fibroblasts (MEFs) and mouse hematopoietic progenitors cells (mHPCs) from WT and FA-A mice. Targeted integration (TI), as well as evidence of hFANCA expression were shown in gene-edited FA-A MEFs. Moreover, evidence of phenotypic correction was demonstrated in these cells by the reversion of the characteristic hypersensitivity to mitomycin C (MMC), and also by the reduction of the MMC-induced chromosomal aberrations. Gene-editing experiments in WT and FA-A mHPCs also allowed us to demonstrate TI in these cell types. As it was observed in FA-A MEFs, evidence of phenotypic correction was also observed in FA-A hematopoietic colonies. In these experiments, we also observed a marked toxicity of plasmid DNA nucleofection that compromised the repopulating properties of gene-edited mHSPCs in irradiated recipients.

### Impact

Our study demonstrates the feasibility of conducting a targeted gene therapy approach in MEFs and HPCs from a mouse model of FA-A and highlights the potential of using for the first time the *Mbs85* locus as a murine *safe harbor* for targeted integration, what opens a new platform that allows the study of the real implication of what means a safe harbor locus in an *in vivo* model prior to the clinic.

## FOR MORE INFORMATION

For more information Fanconi anemia research Foundation: www.anemiadefanconi.org and www.fanconi.org

## EXPANDED VIEW FIGURE LEGENDS

**Expanded View Figure 1. Efficient TALEN-mediated editing of *Mbs85* locus. A)** Schematic representation of the *Mbs85* genomic locus (Chromosome 7: 4,481,520-4,501,680) and the TALEN-targeted sequences (intron 1). The TALEN backbone contains an N-terminal NLS, the ‘0 repeat’, the 17.5 ‘half-repeat’. This is followed bythe C-terminal domain fused to the catalytic *FokI* cleavage domain, as shown in panel B) The amino acids sequence of the DNA binding modules of the engineered TALENs (represented with different colors according to the cipher NG = T, HD = C, NI = A and NN = G or A), as well as the expected target sequences are indicated. **B)** Evaluation of TALEN expression in HEK-293T cells by WB analysis using an antibody directed against the HA-tag present in each TALEN monomer. **C)** Evaluation of the cleavage efficacy of the *Mbs85*-specific TALEN pair using the mismatch sensitive Surveyor assay in WT and FA-A MEFs. Representative electrophoresis gel showing the frequency of indels (calculated as the mean percentage of modified alleles) at the target locus using 2.5 µg of each TALEN monomer. Arrows indicate the size of the parental band (405 bp) and the expected positions of the digestion products (224 bp and 181 bp), that are also indicated with asterisks. φ: transfected cells without DNA (mock condition); IX: DNA molecular weight marker.

**Expanded View Figure 2: Differential repair preference leading to insertions in WT and FA-A MEFs upon nuclease treatment.** Positions of insertions and respective sequences identified at the on-target site. The 24 top most insertions in the on-target locus are shown in **A**) as alignment to the reference sequence, and highlighted in bold and in **B**) as percentage of all insertions comparing WT-MEFs to their respective counterparts FA-A MEFs. Nucleotide positions refer to the amplicon sequence provided in Supplementary Table1.

**Expanded View Figure 3: Differential repair preference leading to deletions in WT and FA-A MEFs upon nuclease treatment.** For all single deletions, the deleted regions and sequences identified at the on-target site. The 24 top most deletions in the on-target locus are shown in a) as alignment to the reference sequence and in b) as percentage of all deletions comparing WT-MEFs to their respective counterparts FA-A MEFs. Percentages of individual deletions were calculated in respect to all deletions. Nucleotide positions refer to the amplicon sequence provided in Supplementary Table1.

**Expanded View Figure 4. On-target cleavage predominantly occurs in the range of nucleotide 222 to 226.** The deletion frequencies of each single nucleotide were calculated as percent of the total reads showing an indel in the analyzed on-target region. Nucleotide positions refer to the amplicon sequence provided in Supplementary Table1.

**Expanded View Figure 5. Evaluation of cleavage efficacy of TALEN in the *Mbs85* locus of WT and FA-A Lin^-^ BM cells by Surveyor assay. A)** Representative electrophoresis gel showing the disruption of the target locus using 2.5 µg of each TALEN monomer indicated as T. The extent of cleavage measured as the mean percentage of modified alleles is indicated below the image. Arrows indicate the size of the parental band (405 bp) and the expected positions of the digestion products (224 bp and 181 bp), that are also indicated with asterisks. U: untransfected cells; IX: DNA molecular weight marker. **B)** Histogram representing the percentage of cleavage monitored with the Surveyor Assay in WT and FA-A hematopoietic progenitors. Bars indicate the mean ± SD (n=3 experiments in WT cells and n=4 experiments in FA-A cells). No statistical differences between groups was observed using a Student’s t-test.

**Expanded View Figure 6. Evaluation of the efficiency of EGFP expression of WT Lin^-^ BM cells nucleofected with the TALEN and the PGK-*EGFP* reporter donor at 2 and 14 days post-nucleofection. A)** Representative flow cytometry dot plots of EGFP^+^ cells analysed at day 2 and 14 post-nucleofection of WT Lin^-^ BM cells. T+D: 0.75 or 2.5 µg of each TALEN monomer together with 4 µg of the PGK-*EGFP* reporter donor; D: 4 µg of the PGK-*EGFP* reporter donor. EGFP^+^ expression was determined discarding autofluorescent cells (576/26 channel). **B)** Representative image in liquid culture of EGFP^+^ Lin^-^ BM cells at 14 days post-nucleofection. EGFP^+^ fluorescent cells could be observed in the microscope in T+D conditions. Scale bar represents 50 μm for all the microphotographs. **C)** Geometric Mean Fluorescent Intensity (measured in arbitrary units, a.u.) of nucleofected cells analysed in the same conditions shown in A). Data show the mean ± S.D. (n=2-3 experiments). Statistical analysis could not be performed.

## EXPANDED VIEW TABLES AND THEIR LEGENDS

**Supplementary Table 1.**
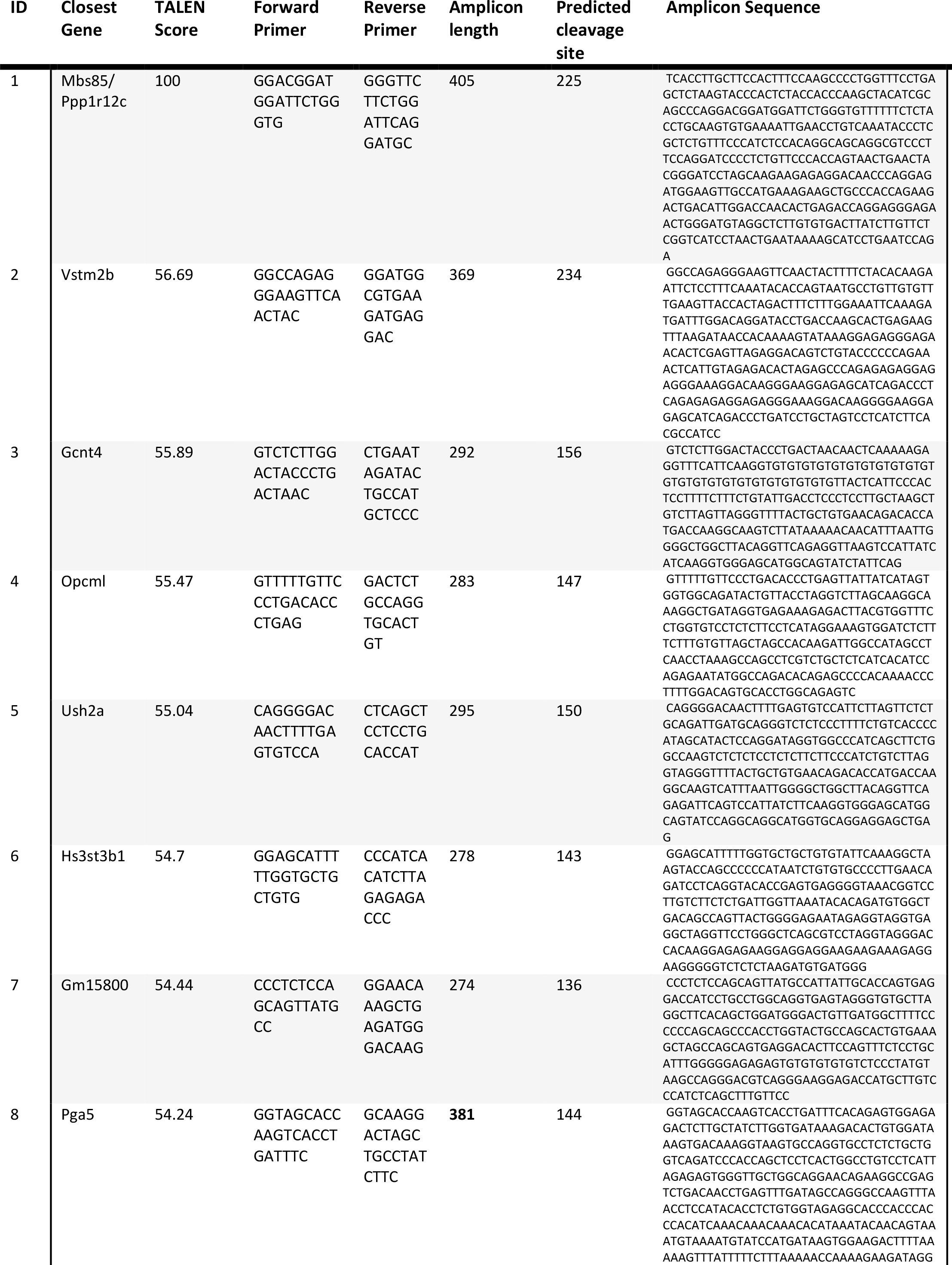
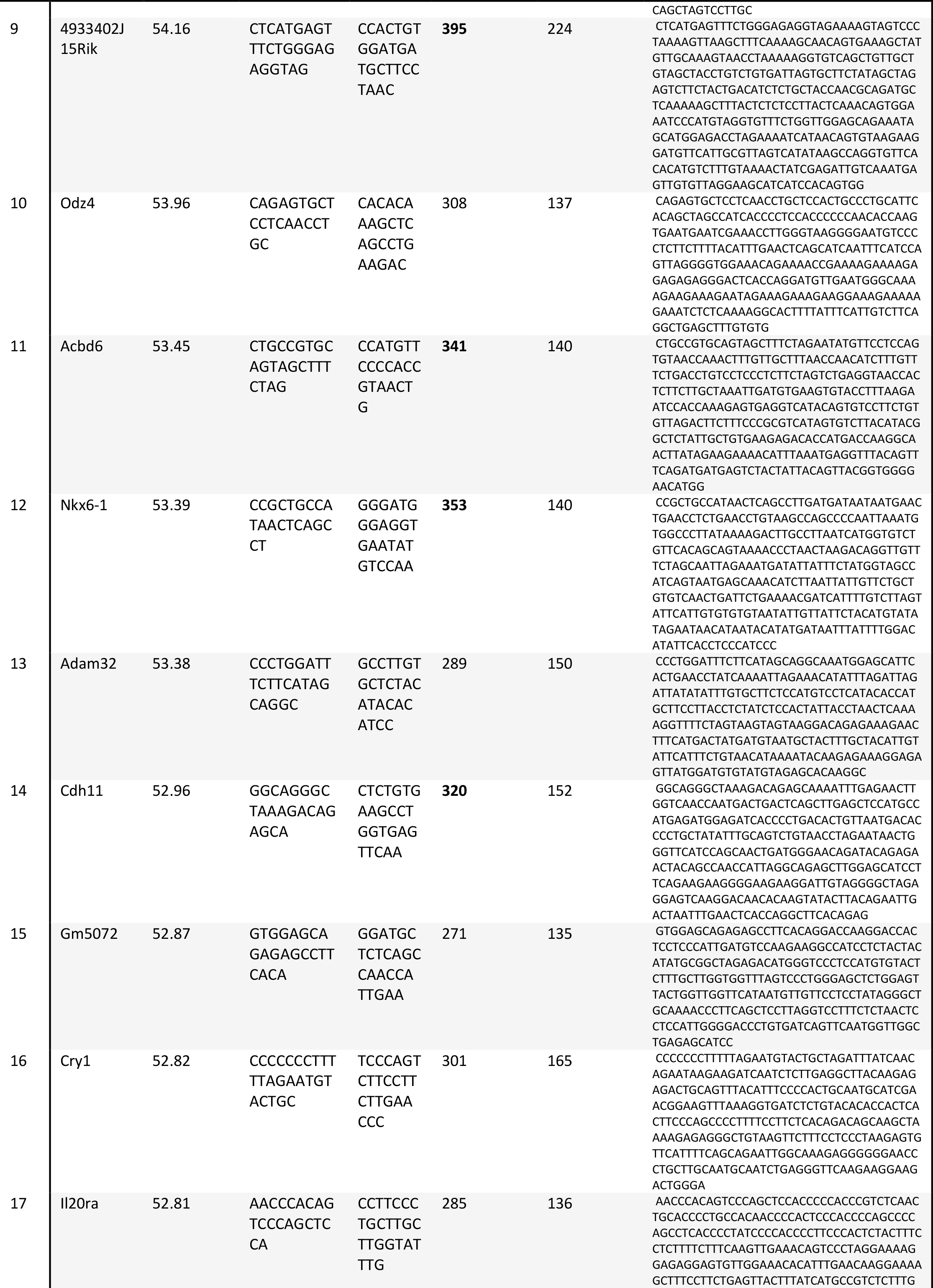
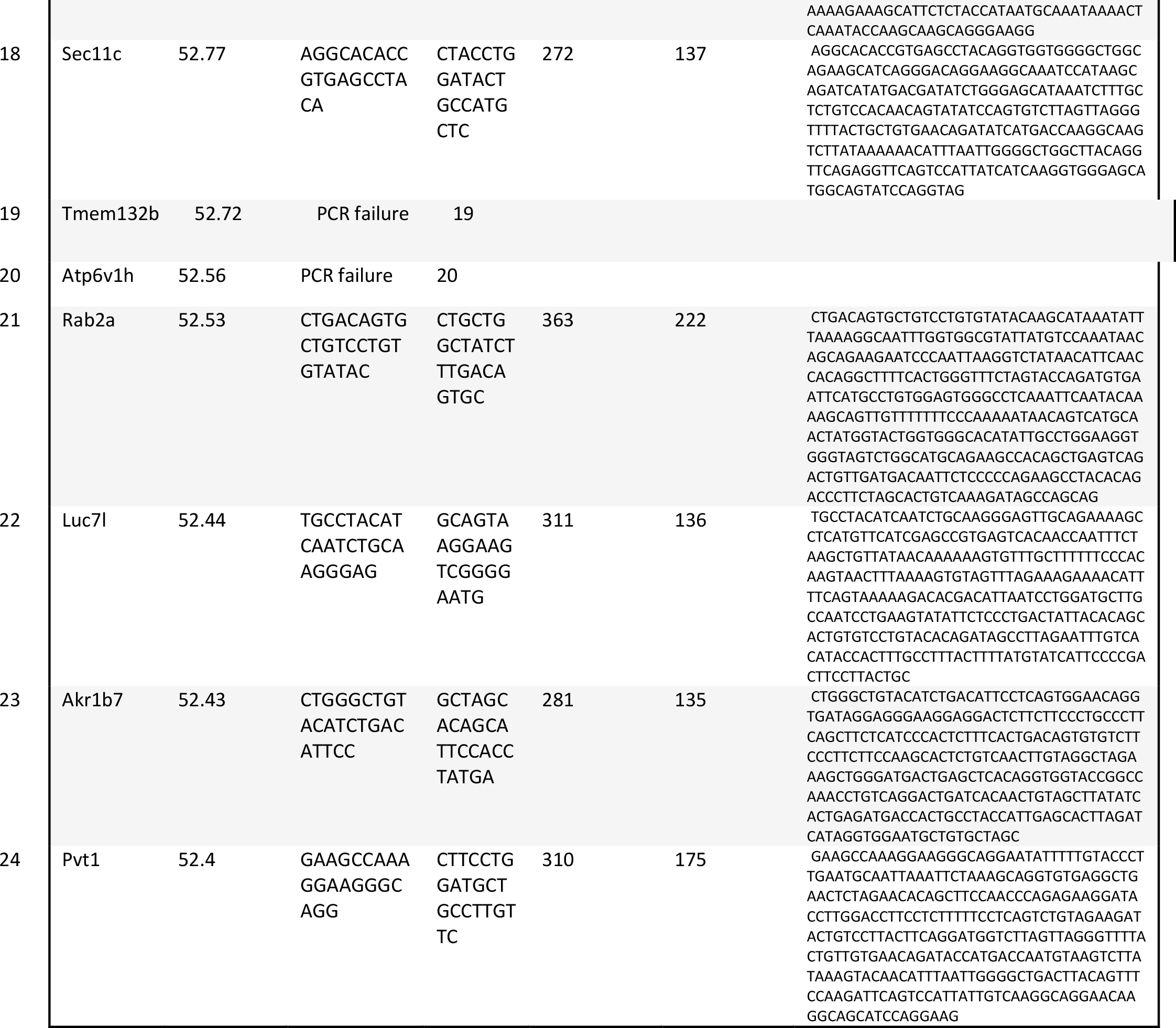
Details of the off-target analyses. The top off-targets are listed according to the ranking generated with TALENv.2 algorithm. The closest gene, the TALEN score, the forward and reverse primers sequence, and the amplicon size and sequence are indicated, as well at the predicted cleavage site.

**Supplementary Table 2.** Results of the next generation sequencing. The top off-targets are listed according to the ranking generated with TALENv.2 algorithm. The frequency of indels generated in each locus could be observed in WT MEFs and their respective counterparts, FA-A MEFs.

| ID | Closest Gene | WT MEFs |  |  |  |  |  | FA-A MEFs |  |  |  |  |  |
| --- | --- | --- | --- | --- | --- | --- | --- | --- | --- | --- | --- | --- | --- |
|  |  | untreated |  |  | nuclease treated |  |  | untreated |  |  | nuclease treated |  |  |
|  |  | Total number of aligned reads | Reads with indels | Indel Frequency | Total number of aligned reads | Reads with indels | Indel Frequency | Total number of aligned reads | Reads with indels | Indel Frequency | Total number of aligned reads | Reads with indels | Indel Frequency |
| 1 | Mbs85/Ppp1r12c | 103938 | 109 | 0.10% | 128806 | 47810 | 37.12% | 107317 | 43 | 0.04% | 97966 | 29534 | 30.15% |
| 2 | Vstm2b | 120329 | 87 | 0.07% | 110015 | 84 | 0.08% | 81881 | 96 | 0.12% | 105707 | 153 | 0.14% |
| 3 | Gcnt4 | 69690 | 74 | 0.11% | 65566 | 44 | 0.07% | 45999 | 37 | 0.08% | 40663 | 26 | 0.06% |
| 4 | Opcml | 91141 | 64 | 0.07% | 55282 | 44 | 0.08% | 117583 | 90 | 0.08% | 95791 | 70 | 0.07% |
| 5 | Ush2a | 100840 | 189 | 0.19% | 78141 | 84 | 0.11% | 90515 | 142 | 0.16% | 113998 | 133 | 0.12% |
| 6 | Hs3st3b1 | 197005 | 134 | 0.07% | 291378 | 205 | 0.07% | 176089 | 120 | 0.07% | 425033 | 338 | 0.08% |
| 7 | Gm15800 | 95634 | 54 | 0.06% | 107987 | 54 | 0.05% | 102713 | 69 | 0.07% | 150677 | 83 | 0.06% |
| 8 | Pga5 | 115423 | 57 | 0.05% | 91053 | 59 | 0.06% | 38234 | 21 | 0.05% | 43245 | 27 | 0.06% |
| 9 | 4933402J15Rik | 74900 | 70 | 0.09% | 72764 | 62 | 0.09% | 84815 | 67 | 0.08% | 75574 | 71 | 0.09% |
| 10 | Odz4 | 195818 | 120 | 0.06% | 201147 | 122 | 0.06% | 200030 | 124 | 0.06% | 185237 | 119 | 0.06% |
| 11 | Acbd6 | 92628 | 79 | 0.09% | 83253 | 79 | 0.09% | 86611 | 70 | 0.08% | 76424 | 72 | 0.09% |
| 12 | Nkx6-1 | 95298 | 141 | 0.15% | 87883 | 107 | 0.12% | 102318 | 166 | 0.16% | 88023 | 119 | 0.14% |
| 13 | Adam32 | 104306 | 93 | 0.09% | 108741 | 396 | 0.36% | 97994 | 108 | 0.11% | 123054 | 403 | 0.33% |
| 14 | Cdh11 | 93757 | 69 | 0.07% | 94248 | 55 | 0.06% | 90752 | 51 | 0.06% | 85379 | 70 | 0.08% |
| 15 | Gm5072 | 156935 | 30349 | 19.34% | 145673 | 26758 | 18.37% | 140998 | 25520 | 18.10% | 142535 | 24233 | 17.00% |
| 16 | Cry1 | 66067 | 94 | 0.14% | 71101 | 101 | 0.14% | 67634 | 89 | 0.13% | 72181 | 95 | 0.13% |
| 17 | Il20ra | 71316 | 57 | 0.08% | 77064 | 81 | 0.11% | 76799 | 93 | 0.12% | 63527 | 46 | 0.07% |
| 18 | Sec11c | 73213 | 48 | 0.07% | 84174 | 47 | 0.06% | 75950 | 56 | 0.07% | 61495 | 47 | 0.08% |
| 19 | Tmem132b | PCR failure |  |  |  |  |  |  |  |  |  |  |  |
| 20 | Atp6v1h | PCR failure |  |  |  |  |  |  |  |  |  |  |  |
| 21 | Rab2a | 63093 | 112 | 0.18% | 81971 | 200 | 0.24% | 69384 | 118 | 0.17% | 64857 | 126 | 0.19% |
| 22 | Luc7l | 178913 | 413 | 0.23% | 196906 | 199 | 0.10% | 235926 | 283 | 0.12% | 188185 | 221 | 0.12% |
| 23 | Akr1b7 | 111556 | 56 | 0.05% | 95369 | 48 | 0.05% | 121969 | 55 | 0.05% | 104876 | 46 | 0.04% |
| 24 | Pvt1 | 112860 | 124 | 0.11% | 113056 | 131 | 0.12% | 116172 | 125 | 0.11% | 111365 | 109 | 0.10% |

**Supplementary Table 3.**
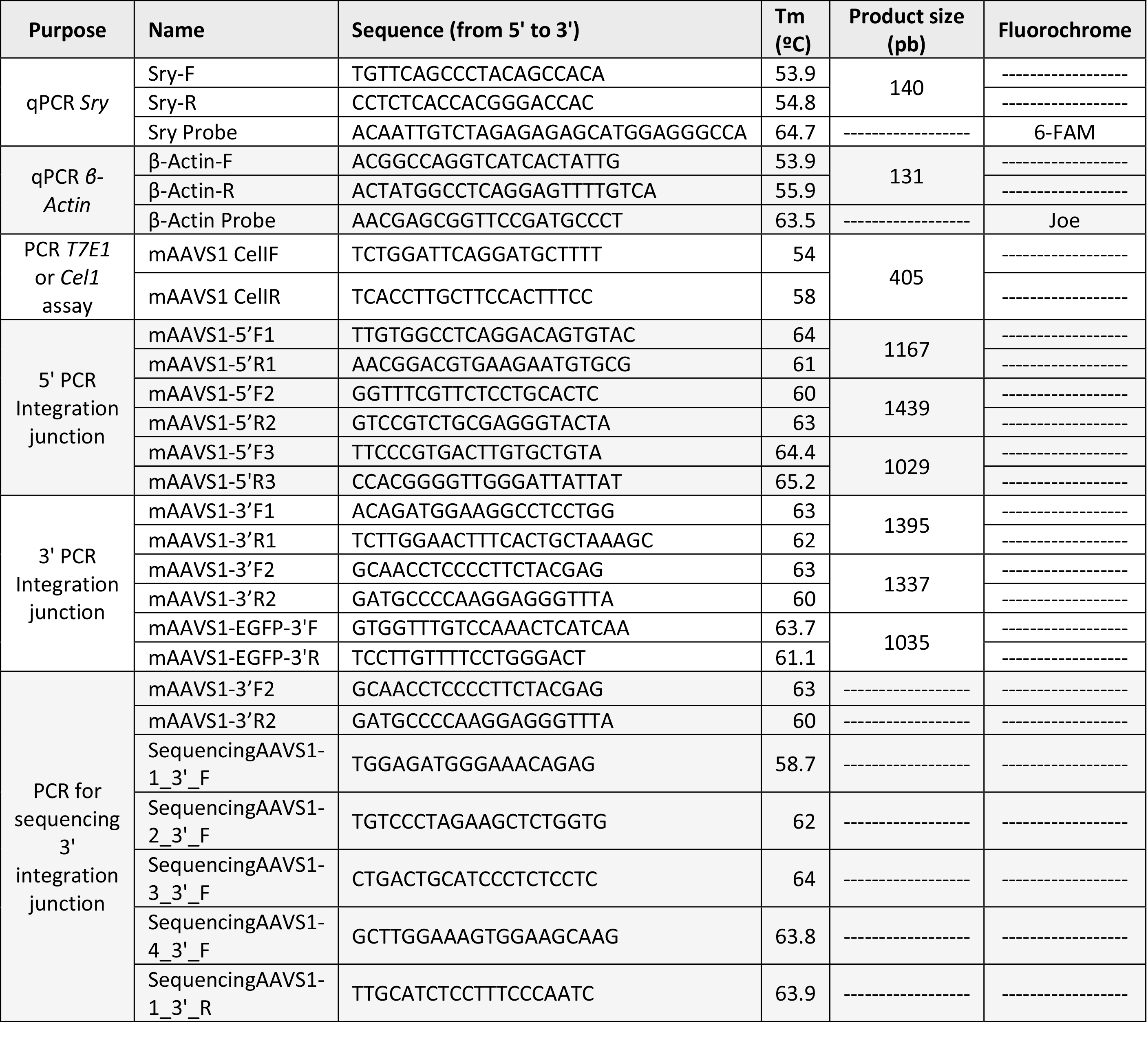
Primers and probes used in this study. Forward and reverse primers as well as probes, their sequence, their melting temperature (Tm) and the product size is indicated. In the case of probes, the fluorochrome to which each probe was conjugated is also indicated.

